# A model for quorum-sensing mediated stochastic biofilm nucleation

**DOI:** 10.1101/2022.03.23.485488

**Authors:** Patrick Sinclair, Chris A. Brackley, Martín Carballo-Pacheco, Rosalind J. Allen

## Abstract

Surface-attached bacterial biofilms cause disease and industrial biofouling, as well as being widespread in the natural environment. Density-dependent quorum sensing is one of the mechanisms implicated in biofilm initiation. Here we present and analyse a model for quorum-sensing triggered biofilm initiation. In our model, individual, planktonic bacteria adhere to a surface, proliferate and undergo a collective transition to a biofilm phenotype. This model predicts a stochastic transition between a loosely attached, finite, layer of bacteria near the surface, and a growing biofilm. The transition is governed by two key parameters: the collective transition density relative to the carrying capacity, and the immigration rate relative to the detachment rate. Biofilm initiation is complex, but our model suggests that stochastic nucleation phenomena may be relevant.

---

Biofilms – surface-attached communities of bacteria surrounded by a polymeric extracellular matrix – are ubiquitous in nature [1]. They play a major role in clinical infections, where they are hard to treat with antibiotics [2], and in industrial biofouling [3]. Therefore, prevention of biofilm initiation is an important clinical and industrial goal. Mathematical models for biofilm growth are well-established [4], ranging from individual-based [5–7], to continuum models [8, 9]. However, most work has focussed on well-established biofilms; few theoretical models exist for early biofilm establishment [10].

The classical picture of biofilm initation starts with reversible attachment of planktonic bacteria to a surface, which may be mediated by flagella and pili. Surface-attached bacteria then proliferate to form microcolonies, and undergo a transition to a different physiological state in which they start to produce the components of the biofilm extracellular matrix, which mediate irreversible attachment to the surface [11]. Regulation of the switch to the biofilm state differs across species. There are many examples of the involvement of collective signalling via quorum sensing [12–15]. However, intracellular cyclic di-GMP signalling, that is not thought to act collectively, also plays a key role [16]. The transition to the biofilm phenotype can also involve biophysical factors such as cell surface motility (which may have collective aspects) [10, 17–19], physical interactions among cells [20] and surface sensing [21].

In this Letter, we present a simple theoretical model for quorum-sensing mediated biofilm initiation. Quorum-sensing describes communication among bacteria that is density-dependent, *i.e.,* it is a form of social interaction [22, 23]. In many bacterial species, extracellular signal molecules called autoinducers (AI) are secreted into the environment; the local concentration of AI provides a way to sense the local bacterial density. Once AI concentration reaches a critical threshold, a gene regulatory response is initiated [22, 23]. This quorum-sensing response can affect a host of different behaviours, including motility, conjugation, competence, sporulation, virulence and biofilm formation.

Using stochastic simulations and simple analytic theory for a coarse-grained micro-habitat model, we investigate the population dynamics of quorum-sensing mediated biofilm initiation. Our model predicts a stochastic transition between a loosely attached, finite, layer of bacteria near the surface, and a growing biofilm. The transition is governed by two key parameters: the threshold population size for the collective transition relative to the carrying capacity, and the immigration rate relative to the detachment rate.

## Microhabitat model

We consider a one-dimensional model in which a growing biofilm is represented as a series of layers, or microhabitats, oriented parallel to the surface (in the *z*-direction; Fig. 1) [4, 24, 25]. The microhabitat has lateral area *a* and depth *δz* = 1 *μ*m (roughly the width of a mono-layer of bacteria). The system initiates with a single, empty microhabitat adjacent to the surface. Bacteria immigrate into this microhabitat from the environment (rate *r*_im_), detach from it (rate *r*_det_), and replicate within it [rate *g*(1 – *N/K*) where *N* is the population size and *K* is the carrying capacity]. To model a quorum-sensing mediated transition to the biofilm phenotype, when the population size reaches a critical value *N*\*, the microhabitat transitions to a “biofilm” state in which bacteria no longer detach. When this happens, a new, adjacent, empty microhabitat is created (Fig. 1). Bacteria can migrate between adjacent microhabitats (rate *r*_mig_). Stochastic simulations of this model were performed using a modified version of Gillespie’s *τ*-leap algorithm [26], in which all events are taken to be Poisson processes [27].

**FIG. 1.**
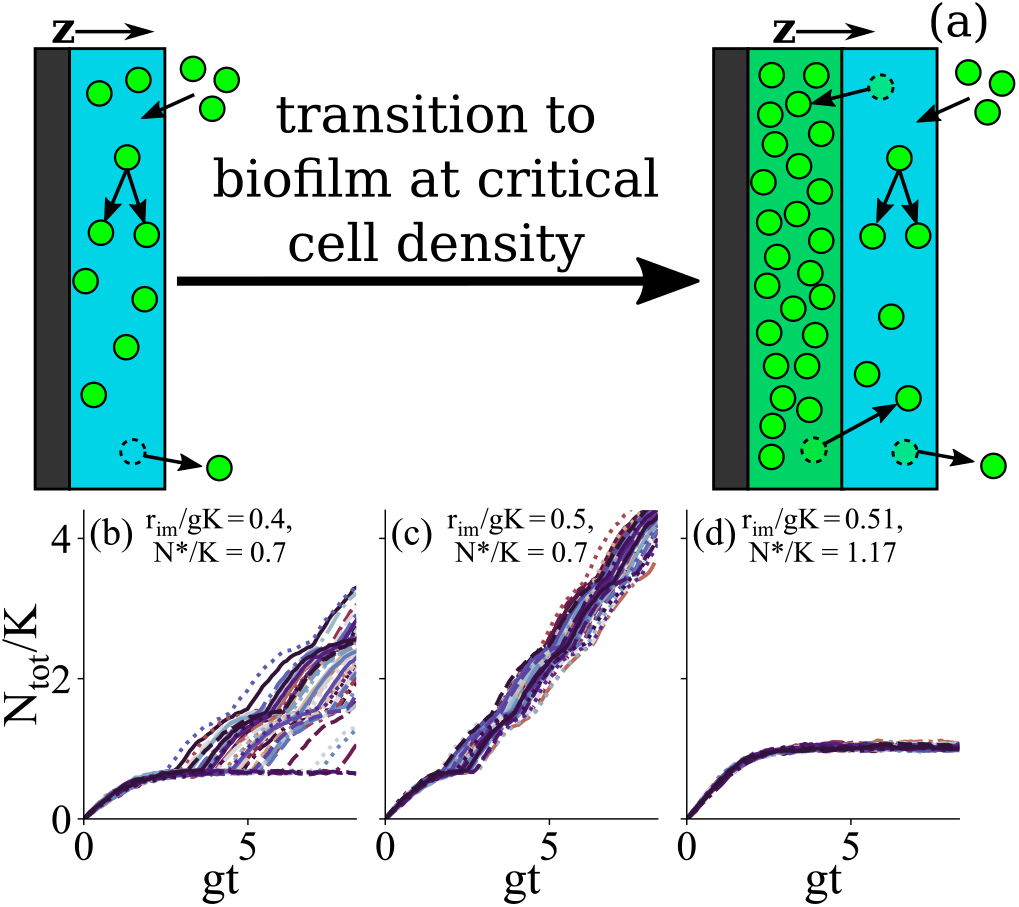
(a) Model for biofilm initiation. The first micro-habitat is shown in blue. Bacteria (green circles) can immigrate, proliferate or detach. Once a critical population size is reached, the first microhabitat transitions to biofilm (green) and a second microhabitat is added. (b)-(d): Stochastic simulation trajectories for different parameter values. The total population size *N*_tot_ is plotted relative to carrying capacity *K*, as a function of dimensionless time *gt*. Other parameter values are *r*_mig_/*g* = 0.8, *r*_det_/*g* = 0.5, *K* = 1000.

In our model, the carrying capacity and the biofilm formation threshold scale with the microhabitat volume (*K* ~ *a δz* and *N*\* ~ *a δz*); the migration rate scales with the microhabitat depth, *r*_mig_ ~ *δz*, and the immigration rate scales with the area, *r*_im_ ~ *a*. Parameters *g* and *r*_det_ are independent of microhabitat dimensions.

Figure 1 shows stochastic trajectories of total population size, for several parameter sets. While the biofilm grows linearly in time from the start in some cases, for other parameters we observe two-step biofilm initiation, in which a “pre-biofilm” forms initially, before it eventually transitions to a growing biofilm. There are also parameter sets for which no biofilm formation is observed for the duration of the simulations.

## Deterministic model

To understand our simulation results, we developed a deterministic version of the model. Considering a system of *M* microhabitats, the number of bacteria in the *i*th microhabitat *N* is described by

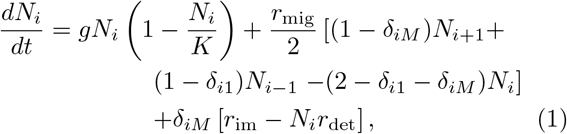

where *δ_ij_* = 1 if *i* = *j* and 0 otherwise. We couple this to the dynamics of increasing numbers of microhabitats where we start with *M* = 1 at *t* = 0 and increase *M* by 1 whenever *N_M_* = *N*\* (*i.e.*, when the outermost, or *M*th microhabitat population reaches the biofilm threshold). Initially *N*_1_ = 0, with *M* = 1.

When *M* = 1, Eqs. (1) reduce to a single equation for *N*_1_. Setting *dN*_1_/*dt* = 0 gives the steady state, and we obtain fixed points

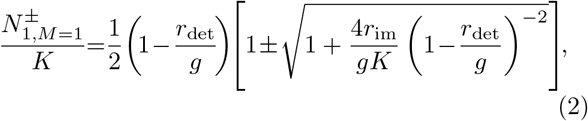

where *M* = 1 in the subscript reminds us that the solution will change once further microhabitats are added. The dimensionless parameter combinations *r*_det_/*g* and *r*_im_/(*Kg*) emerge naturally from Eq. (2). Of these, *r*_det_/*g* is independent of the microhabitat dimensions while *r*_im_/(*Kg*) scales with the microhabitat depth *δz*. Of the two fixed points, the largest one is stable and is always greater than or equal to zero, while the the other one is unstable. Hereon, we will use superscript ‘fp’ to refer to the positive, stable, fixed point.

Two parameter regimes emerge depending on the relative size of the positive fixed point and the biofilm threshold *N*\*. If 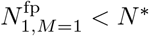, the population in the first microhabitat reaches a steady state [of size 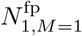 given by the positive, stable solution of Eq. (2)], but the transition to a second microhabitat does not happen and so the biofilm never becomes established. In contrast, if 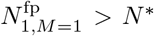, the population in the first microhabitat grows until *N*_1_(*t*) = *N*\*, at which point the first microhabitat transitions to the biofilm state, and a second microhabitat is generated. Fig. 2(a) shows a phase diagram in the (*N*\*, *r*_det_) plane, illustrating the regions of parameter space corresponding to biofilm establishment vs non-establishment (see [27] for further details).

**FIG. 2.**
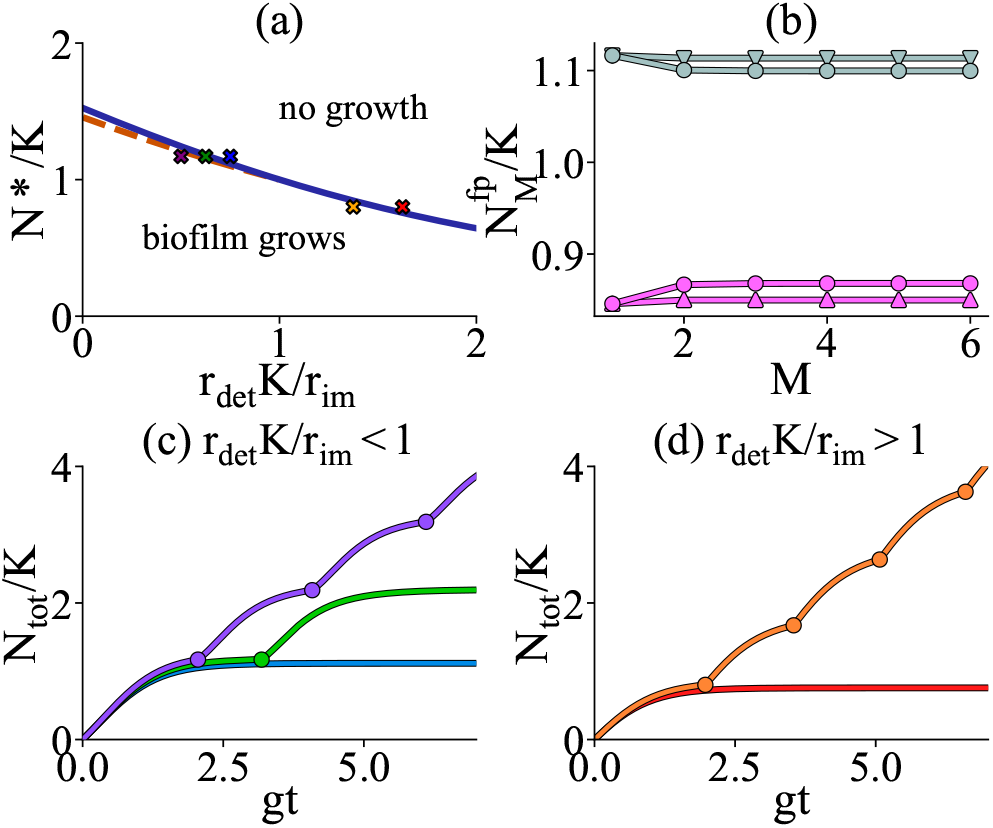
Deterministic model. (a) Phase diagram for *r*_im_/*gK* = 0.8 and *r*_mig_/g = 0.8 (with *K* = 1 and *g* = 1). Solid blue line separates the regime where the population does not reach the biofilm threshold in the first microhabitat and the regime where the biofilm grows. In the region between the dashed and solid blue lines, biofilm will form in the first micro-habitat, but growth will halt in the second. (b) Steady-state population size in the outermost microhabitat for different numbers of microhabitats *M*. Grey points: *r*_det_/*g* = 0.6; pink points: *r*_det_/*g* = 1.1; for round points, *r*_mig_/*g* = 0.8; for triangles, *r*_mig_/*g* = 0.1. In all cases *r*_im_/*gK* = 0.8. Connecting lines are included as a guide for the eye. (c-d) Total population size 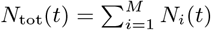 obtained from numerical solution of Eq. (1) with increasing numbers of microhabitats. The parameter values correspond to the coloured crosses in panel (a). In (c) *r*_mig_/*g* = 0.8, *r*_im_/*gK* = 0.8 and *N*/K* = 1.17. From bottom to top the different curves show *r*_det_/*g* = 0.4, 0.5 and 0.6. Points are shown at the times where a new microhabitat is introduced. In (d) *r*_mig_/*g* = 0.8, *r*_im_/*gK* = 0.8 and *N*\* = 0.8. The bottom curve has *r*_det_/*g* = 1.1 and the top curve has *r*_det_/*g* =1.3.

Let us consider further the dynamics in the biofilm establishment regime, 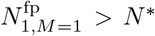. Once the second microhabitat has been generated (*M* = 2), the system is governed by two equations [Eqs. (1) with *i* = 1,2]. To obtain fixed points we now consider *dN*_1_/*dt* = 0, *dN*_2_/*dt* = 0, which leads to two non-linear equations coupled through the migration terms. For a given set of parameters these can be solved numerically. By inspection of the equations it can be inferred that the negative (or zero) fixed point is unstable in favour of the positive one, which we denote 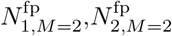 [27]. Interestingly, the population size in the outermost microhabitat 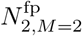 can be larger or smaller than 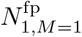 depending on the value of the ratio *r*_det_ *K/r*_im_, which measures the relative importance of detachment and immigration in the outer microhabitat. If detachment dominates, migration tends to be from the inner to the outer microhabitat, whereas if immigration dominates, they tend to migrate from the outer to the inner microhabitat. Thus, as the biofilm grows, its outer edge may become either more or less dense (Fig. 2(b); see also [27]).

In the immigration-dominated regime where *r*_det_*K/r*_im_ < 1, the value of the fixed point in the *M*^th^ microhabitat increases with *M* [Fig. 2(b)]. Therefore there exists a small region of parameter space where it is possible to choose *N*\* such that 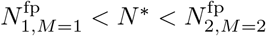 [the region between the dashed and solid lines in Fig. 2(a)]. For such parameters, the first microhabitat transitions to biofilm but the second microhabitat does not, *i.e.,* the system forms a biofilm of finite width. We note that this phenomenon occurs for *N*\* > *K* [Fig. 2(a)], i.e. when the environment does not sustain a high enough steady state population density to trigger biofilm formation. In the detachment-dominated regime where *r*_det_*K/r*_im_ > 1, the biofilm will always show sustained growth if 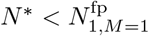.

To study the dynamics of this sustained growth regime we integrate Eqs. (1) numerically. We can obtain population trajectories by starting with the *i* =1 equation, integrating until *N*_1_ = *N*\*, adding the *i* = 2 equation, continuing to integrate both equations until *N*_2_ = *N*\*, and so on. Figs. 2(c-d) show trajectories for the total population size obtained for different parameters (a full exploration of the effect of different parameters is given in [27]). As discussed above, for *r*_det_*K/r*_im_ < 1, depending on the value of N* relative to the fixed points, either the biofilm will never establish, it will establish but be limited to a finite size, or it will continue to grow indefinitely. For *r*_det_ *K/r*_im_ > 1 there are only two possible regimes – non-establishment or continued growth.

## Role of stochasticity

Our deterministic analysis can help us better understand our stochastic simulations [Figs. 1(b-d)]. First, the existence of the different growth regimes explains why for some parameters our stochastic system grows, but for others it does not. Fig. 3 shows a heat-map from our stochastic simulations of the time taken for the system to form biofilm (i.e., for the first microhabitat to exceed the transition threshold *N*\*) as a function of the parameter combinations *N*\*/*K* and *r*_det_*K/r*_im_. Where no colour is shown, the simulations did not form biofilm. The dashed pink line shows the prediction from the deterministic theory for the boundary between biofilm initiation and non-initiation. Strikingly, Fig. 3 shows stochastic biofilm growth in the region of parameter space beyond the deterministic phase boundary.

**FIG. 3.**
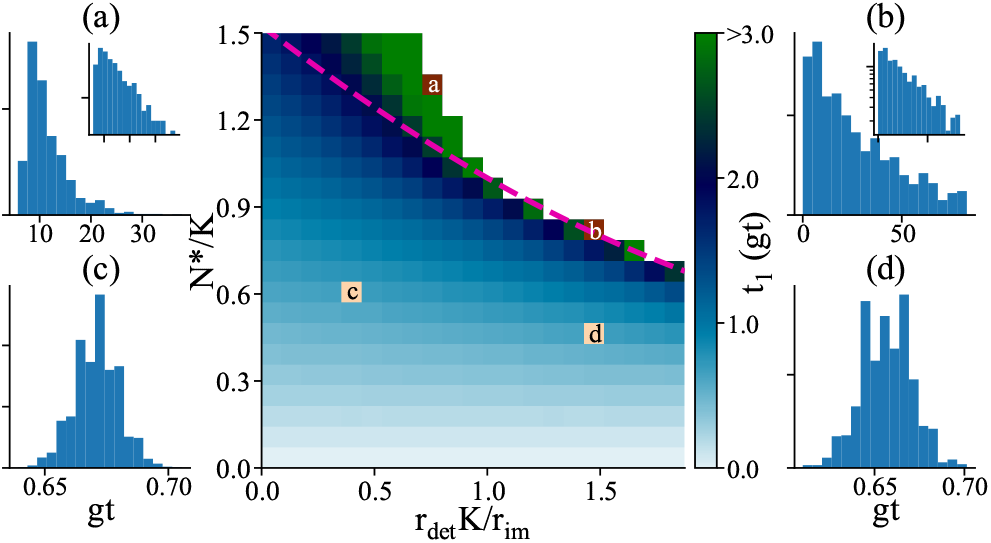
Stochastic initiation dynamics. Centre: Heatmap showing lag time, *t*_1_, before biofilm initiation (*i.e.,* until the first microhabitat reaches *N*\*) in our stochastic simulations, as a function of *N*/K* and *r*_det_*K/r*_im_. Here *r*_im_/*gK* = 0.8 and *r*_mig_/*g* = 0.8, with *K* = 10000 and *g* = 0.083, while *r*_det_ and *N*\* were varied. We use dimensionless time *gt.* The white region in the upper right quadrant denotes parameters where the biofilm does not initiate within the maximum simulation time of ~83*gt*. The dashed pink line shows the deterministic phase boundary [as in Fig. 2(a)]. Panels (a-d) show lag time distributions for the corresponding parameter sets indicated in the heatmap. Distributions (a) and (b) are close to exponential (insets show the same plots on a log scale) with coefficients of variation (CV) 0.367 and 0.807 respectively. Distributions (c) and (d) are closer to Gaussian, with CV values of 0.014 and 0.021.

To understand this, we compare trajectories of biofilm growth from our stochastic simulations to those predicted by the deterministic model (Fig. 4). In the region of parameter space where growth is predicted deterministically, the biofilm grows from the start, and the deterministic and stochastic simulation results match quantitatively [Figs. 4(a,b)]. However for some parameter sets where there is no growth in the deterministic model the stochastic simulations show a lag followed by a transition to growth [Figs. 4(c,d)]. The lag time varies between replicate simulations, with parameter values which lead to a longer mean lag time also showing more lag time variability [compare blue and green lines in Fig. 4(c)]. Indeed, the distribution of lag times before the threshold *N*\* is reached is close to exponential for these simulations [Figs. 3(a,b)], suggesting an underlying Poisson process. In contrast, the waiting time distribution for biofilm formation is narrow and approximately Gaussian for parameter sets in the deterministic growth regime [Figs. 3(c,d)].

**FIG. 4.**
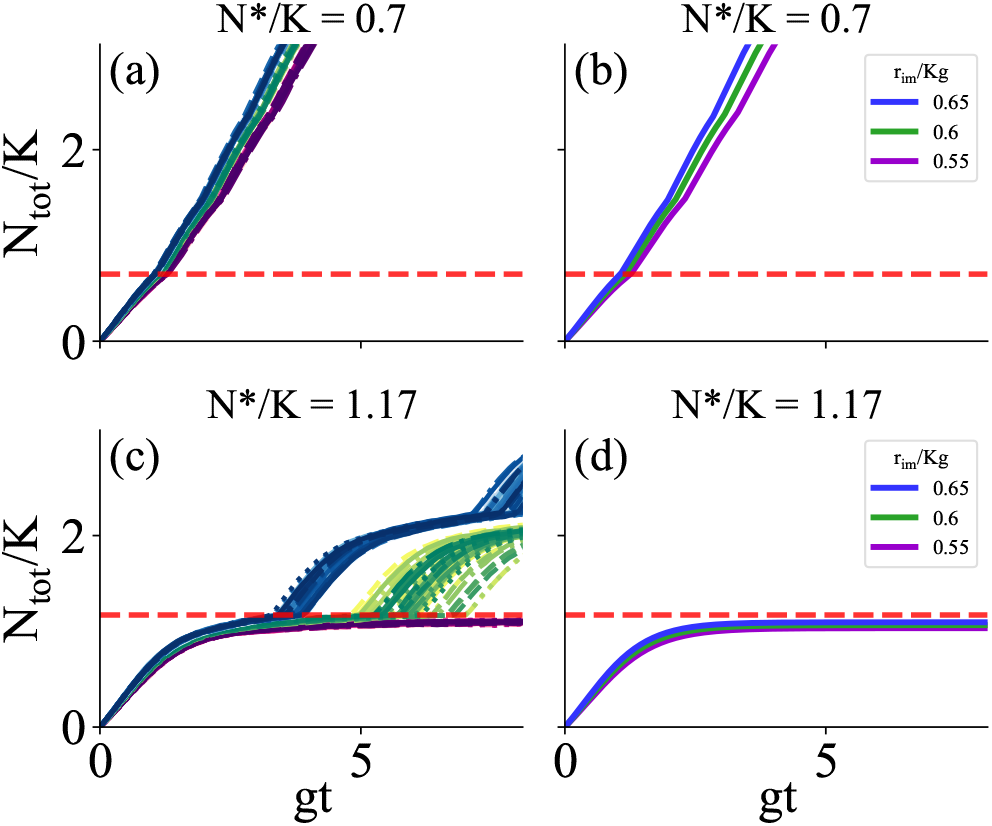
Comparison between stochastic (left-hand panels) and deterministic (right-hand panels) biofilm growth trajectories. (a,b) Parameters in the deterministic growth regime; *N*/K* = 0.7. (c,d) Parameters in the stochastic growth regime; *N*/K* = 1.17. In all cases, *r*_mig_/*g* = 0.8, *r*_det_/*g* = 0.5, *K* = 10,000 and the immigration rate is varied: *r*_im_/*gK* = 0.55, 0.6 and 0.65. The red dashed lines indicate the biofilm transition threshold *N*/K*.

Our results point to the following scenario. For parameters where the deterministic fixed point population size 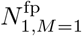 in the first microhabitat is only slightly smaller than the threshold *N*\*, the system will grow to the fixed point, and then a small fluctuation in population size can push the system over the threshold, generating a second microhabitat. The system displays “one-way” dynamics: once new microhabitats form, they cannot be removed. This then amounts to a first passage time process, in which growth, immigration and detachment all play a role. If 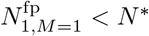, we have a first passage stochastic process with a metastable potential minimum at 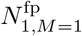, and the lag time observed in Figs. 3(a,b) is the time to escape this minimum and reach the absorbing state at *N*\*.

Consistent with this picture, biofilm initiation can be suppressed in our model by increasing the system size, and hence decreasing the relative magnitude of stochastic fluctuations.Simulations for larger values of the lateral area *a* (which scales *K, N*\* and *r*_im_) show much longer mean lag times in the stochastic growth regime, or no biofilm initiation [27].

In this letter we have presented a coarse-grained micro-habitat model for quorum-sensing mediated biofilm initiation, where a transition to the biofilm state happens when the population density close to the surface crosses a critical threshold. The model accounts for immigration, detachment, proliferation and migration within the biofilm (once several microhabitats have formed). Our work points to different modes of biofilm initiation under different parameter regimes. Under favourable conditions, biofilm growth can initiate immediately. However under less favourable conditions, a loosely attached layer of bacteria first forms at the surface, which may or may not undergo a stochastic transition to biofilm growth.

In our model, the boundary between deterministic and stochastic biofilm initiation depends on whether the biofilm threshold *N*\* is smaller or larger than the deterministic steady-state population size. Quorum sensing is generally studied in liquid cultures, where quorum-sensing regulated genes become activated at cell densities of ~10^9^ml^-1^, although this can vary [28]. For surfaces that can sustain a densely-packed monolayer of cells (*e.g.,* nutrient agar), the steady-state population density is clearly well above this quorum sensing threshold. However this may not be the case in, for example, a marine environment, where nutrient availability is poor. Therefore we might predict greater stochasticity in biofilm initiation under conditions of poor nutrient (small K). Indeed, understanding biofilm initiation is particularly important in marine environments, where biofouling on ship hulls and marine installations is a major concern [29].

Previous theoretical models for biofilm initiation have focused on *Pseudomonas aeruginosa,* where motile cells explore the surface before committing to surface attachment via the production of exopolysaccharides (EPS). The motile cells can leave trails of EPS that affect other cells [10, 18], leading to collective phenomena. Although these models are conceptually very different from ours, both involve collective effects, and a transition from reversible to irreversible attachment.

Our model is of course highly idealised: quorum sensing may not be involved in biofilm initiation for all microbes, and even where it is, the process is far more complex than our representation (e.g., a biofilm may even initiate under conditions of low rather than high density [13, 14]). Our use of discrete microhabitats, while conceptually simple and computationally efficient, omits important biological information, such as the configuration and motility of individual bacteria on the surface [18, 20], aggregation prior to biofilm formation [30, 31] and the multispecies nature of many environmental biofilms. Nevertheless, our work suggests that biofilm initiation can be an intrinsically stochastic process – with potentially important implications for our ability to predict and control biofilm infections and industrial biofouling processes.

Laboratory biofilm growth experiments are notoriously difficult to reproduce quantitatively, and some studies have quantified biofilm growth variability [32, 33]. However, such quantification is challenging, because well-controlled, quantitative biofilm growth experiments can be technically difficult, due to inevitable spatial heterogeneity within flow devices, feedback between biofilm growth and flow patterns, and differences in the state of the inoculating microbes, among other factors [33]. Our work suggests that, although difficult, investigation of intrinsic biofilm stochasticity may prove interesting, fruitful, and relevant.

PS was supported by an EPSRC NPIF studentship, and RJA and MCP were funded by the European Research Council under Consolidator grant 682237 EVOSTRUC. RJA was also supported by the Excellence Cluster Balance of the Microverse (EXC 2051 - Project-ID 390713860) funded by the Deutsche Forschungsgemeinschaft (DFG). CAB was funded by the European Research Council under Consolidator Grant 648050 THREEDCELLPHYSICS. The authors thank Jennifer Longyear, Kevin Reynolds, Clayton Price, Nick Cogan and Paul Hush for valuable discussions.

## References

[1] H.-C. Flemming and S. Wuertz, Nat. Rev. Microbiol. 17, 247 (2019).

[2] T.-F. C. Mah and G. A. O’Toole, Trends Microbiol. 9, 34 (2001).

[3] P. S. Murthy and R. Venkatesan, “Industrial biofilms and their control,” in Marine and Industrial Biofouling, edited by H.-C. Flemming, P. S. Murthy, R. Venkatesan, and K. Cooksey (Springer Berlin Heidelberg, Berlin, Heidelberg, 2009) pp. 65–101.

[4] R. J. Allen and B. a. Waclaw, Rep. Prog. Phys. 82, 016601 (2019).

[5] J. B. Xavier, C. Picioreanu, and M. C. M. van Loosdrecht, Environmental Microbiology 7, 1085 (2005).

[6] L. A. Lardon, B. V. Merkey, S. Martins, A. Dötsch, C. Picioreanu, J. U. Kreft, and B. F. Smets, Environmental Microbiology 13, 2416 (2011).

[7] B. Li, D. Taniguchi, J. P. Gedara, V. Gogulancea, R. Gonzalez-Cabaleiro, J. Chen, A. S. McGough, I. D. Ofiteru, T. P. Curtis, and P. Zuliani, PLoS Comput. Biol. 15, e1007125 (2019).

[8] T. Zhang, N. Cogan, and Q. Wang, Commun. Comput. Phys. 4, 72 (2008).

[9] T. Zhang, N. Cogan, and Q. Wang, SIAM J. Appl. Math 69, 641 (2008).

[10] A. Gelimson, K. Zhao, C. K. Lee, W. T. Kranz, G. C. L. Wong, and R. Golestanian, Phys. Rev. Lett. 117, 178102 (2016).

[11] C. Berne, C. K. Ellison, A. Ducret, and Y. V. Brun, Nat. Rev. Microbiol. 16, 616 (2018).

[12] D. G. Davies, M. R. Parsek, J. P. Pearson, B. H. Iglewski, J. W. Costerton, and E. P. Greenberg, Science 280, 295 (1998).

[13] B. K. Hammer and B. L. Bassler, Mol. Microbiol. 50, 101 (2003).

[14] J. M. Yarwood, D. J. Bartels, E. M. Volper, and P. Greenberg, J. Bacteriol. 186, 1838 (2004).

[15] M. D. Koutsoudis, D. Tsaltas, T. D. Minogue, and S. B. von Bodman, Proc. Natl. Acad. Sci. USA 103, 5983 (2006).

[16] M. Valentini and A. Filloux, J. Biol. Chem. 291, 12547 (2016).

[17] M. L. Gibiansky, J. C. Conrad, F. Jin, V. D. Gordon, D. A. Motto, M. A. Mathewson, W. G. Stopka, D. C. Zelasko, J. D. Shrout, and G. C. L. Wong, Science 330, 197 (2010).

[18] K. Zhao, B. S. Tseng, B. B., F. Jin, M. L. Gibiansky, J. J. Harrison, E. Luijten, M. R. Parsek, and G. C. L. Wong, Nature 497, 338 (2013).

[19] C. K. Lee, J. Vachier, J. de Anda, K. Zhao, A. E. Baker, R. R. Bennett, C. R. Armbruster, K. A. Lewis, T. R. L., C. J. Lomba, D. A. Hogan, M. R. Parsek, G. A. O’Toole, R. Golestanian, and G. C. L. Wong, mBio 11, e02644 (2020).

[20] F. Beroz, J. Yan, Y. Meir, B. Sabass, H. A. Stone, B. L. Bassler, and N. S. Wingreen, Nat. Phys. 14, 954 (2018).

[21] G. A. O’Toole and G. C. L. Wong, Curr. Opin. Microbiol. 30, 139 (2016).

[22] C. M. Waters and B. L. Bassler, Annu. Rev. Cell Dev. Biol. 21, 319 (2005).

[23] M. B. Miller and B. L. Bassler, Annu. Rev. Microbiol. 55, 165 (2001).

[24] P. Greulich, B. Waclaw, and R. J. Allen, Physical Review Letters 109, 088101 (2012).

[25] P. Sinclair, M. Carballo-Pacheco, and R. J. Allen, Phys. Biol. 16, 046001 (2019).

[26] Y. Cao, D. T. Gillespie, and L. R. Petzold, J. Chem. Phys. 123, 054104 (2005).

[27] See the online Supplementary Material.

[28] R. Siehbel, B. Traxler, D. D. An, M. R. Parsek, A. L. Schaefer, and P. K. Singh, Proc. Natl. Acad. Sci. U. S. A. 107, 7916 (2010).

[29] T. R. Bott, Industrial biofouling (Elsevier, 2011).

[30] K. N. Kragh, J. B. Hutchison, G. Melaugh, C. Rodesney, A. E. L. Roberts, Y. Irie, P. Jensen, S. P. Diggle, R. J. Allen, V. Gordon, and T. Bjarnsholt, mBio 7, e00237 (2016).

[31] G. Melaugh, J. Hutchison, K. N. Kragh, Y. Irie, A. Roberts, T. Bjarnsholt, S. P. Diggle, V. D. Gordon, and R. J. Allen, PLoS One 11, e0149683 (2016).

[32] O. Kroukamp, R. G. Dumitrache, and G. M. Wolfaardt, Appl. Env. Microbiol. 76, 6025 (2010).

[33] A. Heydorn, K. Ersboll, B, M. Hentzer, M. R. Parsek, M. Givskov, and S. Molin, Microbiology 146, 2409 (2000).

