## Supplemental Material for "A model for quorum-sensing mediated stochastic biofilm nucleation"

In our model, the carrying capacity and the biofilm formation threshold scale with the microhabitat volume ($K \sim a\,\delta z$ and $N^* \sim a\,\delta z$); the migration rate scales with the microhabitat depth, $r_{\mathrm{mig}} \sim \delta z$, and the immigration

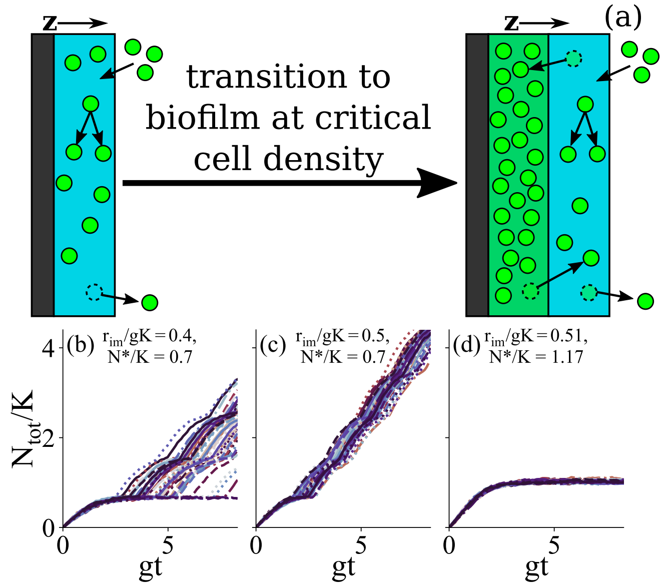


FIG. 1. (a) Model for biofilm initiation. The first microhabitat is shown in blue. Bacteria (green circles) can immigrate, proliferate or detach. Once a critical population size is reached, the first microhabitat transitions to biofilm (green) and a second microhabitat is added. (b)-(d): Stochastic simulation trajectories for different parameter values. The total population size $N_{\rm tot}$ is plotted relative to carrying capacity $K$, as a function of dimensionless time $gt$. Other parameter values are $r_{\rm mig}/g = 0.8$, $r_{\rm det}/g = 0.5$, $K = 1000$.

rate scales with the area, $r_{\rm im} \sim a$. Parameters $g$ and $r_{\rm det}$ are independent of microhabitat dimensions.

Figure 1 shows stochastic trajectories of total population size, for several parameter sets. While the biofilm grows linearly in time from the start in some cases, for other parameters we observe two-step biofilm initiation, in which a "pre-biofilm" forms initially, before it eventually transitions to a growing biofilm. There are also parameter sets for which no biofilm formation is observed for the duration of the simulations.

*Deterministic model.*—To understand our simulation results, we developed a deterministic version of the model. Considering a system of $M$ microhabitats, the number of bacteria in the $i$th microhabitat $N_i$ is described by

$$\frac{dN_i}{dt} = gN_i\left(1-\frac{N_i}{K}\right) + \frac{r_{\rm mig}}{2}\left[(1-\delta_{iM})N_{i+1} + (1-\delta_{i1})N_{i-1} - (2-\delta_{i1}-\delta_{iM})N_i\right] + \delta_{iM}\left[r_{\rm im} - N_i r_{\rm det}\right], \qquad (1)$$

where $\delta_{ij} = 1$ if $i = j$ and 0 otherwise. We couple this to the dynamics of increasing numbers of microhabitats where we start with $M = 1$ at $t = 0$ and increase $M$ by 1 whenever $N_M = N^*$ (*i.e.*, when the outermost, or $M$th microhabitat population reaches the biofilm threshold). Initially $N_1 = 0$, with $M = 1$.

When $M = 1$, Eqs. (1) reduce to a single equation for $N_1$. Setting $dN_1/dt = 0$ gives the steady state, and we obtain fixed points

$$\frac{N^{\pm}_{1,M=1}}{K} = \frac{1}{2}\left(1-\frac{r_{\rm det}}{g}\right)\left[1 \pm \sqrt{1+\frac{4r_{\rm im}}{gK}\left(1-\frac{r_{\rm det}}{g}\right)^{-2}}\,\right], \qquad (2)$$

where $M = 1$ in the subscript reminds us that the solution will change once further microhabitats are added. The dimensionless parameter combinations $r_{\rm det}/g$ and $r_{\rm im}/(Kg)$ emerge naturally from Eq. (2). Of these, $r_{\rm det}/g$ is independent of the microhabitat dimensions while $r_{\rm im}/(Kg)$ scales with the microhabitat depth $\delta z$. Of the two fixed points, the largest one is stable and is always greater than or equal to zero, while the the other one is unstable. Hereon, we will use superscript 'fp' to refer to the positive, stable, fixed point.

Two parameter regimes emerge depending on the relative size of the positive fixed point and the biofilm threshold $N^*$. If $N^{\rm fp}_{1,M=1} < N^*$, the population in the first microhabitat reaches a steady state [of size $N^{\rm fp}_{1,M=1}$ given by the positive, stable solution of Eq. (2)], but the transition to a second microhabitat does not happen and so the biofilm never becomes established. In contrast, if $N^{\rm fp}_{1,M=1} > N^*$, the population in the first microhabitat grows until $N_1(t) = N^*$, at which point the first microhabitat transitions to the biofilm state, and a second microhabitat is generated. Fig. 2(a) shows a phase diagram in the ($N^*$, $r_{\rm det}$) plane, illustrating the regions of parameter space corresponding to biofilm establishment vs non-establishment (see [27] for further details).

Let us consider further the dynamics in the biofilm establishment regime, $N^{\rm fp}_{1,M=1} > N^*$. Once the second microhabitat has been generated ($M = 2$), the system is governed by two equations [Eqs. (1) with $i = 1, 2$]. To obtain fixed points we now consider $dN_1/dt = 0$, $dN_2/dt = 0$, which leads to two non-linear equations coupled through the migration terms. For a given set of parameters these can be solved numerically. By inspection of the equations it can be inferred that the negative (or zero) fixed point is unstable in favour of the positive one, which we denote $N^{\rm fp}_{1,M=2}$,$N^{\rm fp}_{2,M=2}$ [27]. Interestingly, the population size in the outermost microhabitat $N^{\rm fp}_{2,M=2}$ can be larger or smaller than $N^{\rm fp}_{1,M=1}$ depending on the value of the ratio $r_{\rm det}K/r_{\rm im}$, which measures the relative importance of detachment and immigration in the outer microhabitat. If detachment dominates, migration tends to be from the inner to the outer microhabitat, whereas if immigration dominates, they tend to migrate from the outer to the inner microhabitat. Thus, as the biofilm grows, its outer edge may become either more or less dense (Fig. 2(b); see also [27]).

In the immigration-dominated regime where $r_{\rm det}K/r_{\rm im} < 1$, the value of the fixed point in the

$M^{\text{th}}$ microhabitat increases with $M$ [Fig. 2(b)]. Therefore there exists a small region of parameter space where it is possible to choose $N^*$ such that $N^{\text{fp}}_{1,M=1} < N^* < N^{\text{fp}}_{2,M=2}$ [the region between the dashed and solid lines in Fig. 2(a)]. For such parameters, the first microhabitat transitions to biofilm but the second microhabitat does not, *i.e.*, the system forms a biofilm of finite width. We note that this phenomenon occurs for $N^* > K$ [Fig. 2(a)], i.e. when the environment does not sustain a high enough steady state population density to trigger biofilm formation. In the detachment-dominated regime where $r_{\text{det}}K/r_{\text{im}} > 1$, the biofilm will always show sustained growth if $N^* < N^{\text{fp}}_{1,M=1}$.

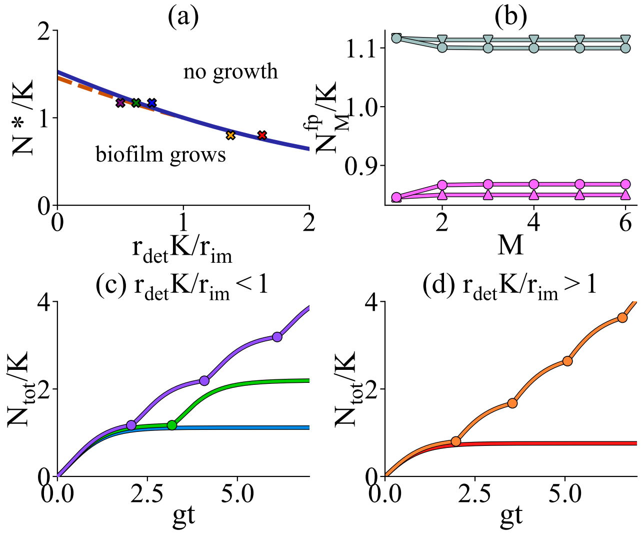


FIG. 2. Deterministic model. (a) Phase diagram for $r_{\text{im}}/gK = 0.8$ and $r_{\text{mig}}/g = 0.8$ (with $K = 1$ and $g = 1$). Solid blue line separates the regime where the population does not reach the biofilm threshold in the first microhabitat and the regime where the biofilm grows. In the region between the dashed and solid blue lines, biofilm will form in the first microhabitat, but growth will halt in the second. (b) Steady-state population size in the outermost microhabitat for different numbers of microhabitats $M$. Grey points: $r_{\text{det}}/g = 0.6$; pink points: $r_{\text{det}}/g = 1.1$; for round points, $r_{\text{mig}}/g = 0.8$; for triangles, $r_{\text{mig}}/g = 0.1$. In all cases $r_{\text{im}}/gK = 0.8$. Connecting lines are included as a guide for the eye. (c-d) Total population size $N_{\text{tot}}(t) = \sum_{i=1}^{M} N_i(t)$ obtained from numerical solution of Eq. (1) with increasing numbers of microhabitats. The parameter values correspond to the coloured crosses in panel (a). In (c) $r_{\text{mig}}/g = 0.8$, $r_{\text{im}}/gK = 0.8$ and $N^*/K = 1.17$. From bottom to top the different curves show $r_{\text{det}}/g = 0.4$, 0.5 and 0.6. Points are shown at the times where a new microhabitat is introduced. In (d) $r_{\text{mig}}/g = 0.8$, $r_{\text{im}}/gK = 0.8$ and $N^* = 0.8$. The bottom curve has $r_{\text{det}}/g = 1.1$ and the top curve has $r_{\text{det}}/g = 1.3$.

Indeed, the distribution of lag times before the threshold $N^*$ is reached is close to exponential for these simulations [Figs. 3(a,b)], suggesting an underlying Poisson process. In contrast, the waiting time distribution for biofilm formation is narrow and approximately Gaussian for parameter sets in the deterministic growth regime [Figs. 3(c,d)].

Our results point to the following scenario. For parameters where the deterministic fixed point population size $N^{\text{fp}}_{1,M=1}$ in the first microhabitat is only slightly smaller than the threshold $N^*$, the system will grow to the fixed point, and then a small fluctuation in population size can push the system over the threshold, generating a second microhabitat. The system displays "one-way" dynamics: once new microhabitats form, they cannot be removed. This then amounts to a first passage time process, in which growth, immigration and detachment all play a role. If $N^{\text{fp}}_{1,M=1} < N^*$, we have a first passage stochastic

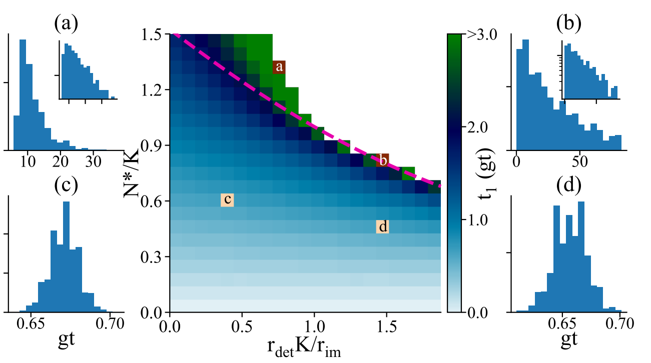


FIG. 3. Stochastic initiation dynamics. Centre: Heatmap showing lag time, $t_1$, before biofilm initiation (*i.e.*, until the first microhabitat reaches $N^*$) in our stochastic simulations, as a function of $N^*/K$ and $r_{\rm det}K/r_{\rm im}$. Here $r_{\rm im}/gK = 0.8$ and $r_{\rm mig}/g = 0.8$, with $K = 10000$ and $g = 0.083$, while $r_{\rm det}$ and $N^*$ were varied. We use dimensionless time $gt$. The white region in the upper right quadrant denotes parameters where the biofilm does not initiate within the maximum simulation time of $\sim$83$gt$. The dashed pink line shows the deterministic phase boundary [as in Fig. 2(a)]. Panels (a-d) show lag time distributions for the corresponding parameter sets indicated in the heatmap. Distributions (a) and (b) are close to exponential (insets show the same plots on a log scale) with coefficients of variation (CV) 0.367 and 0.807 respectively. Distributions (c) and (d) are closer to Gaussian, with CV values of 0.014 and 0.021.

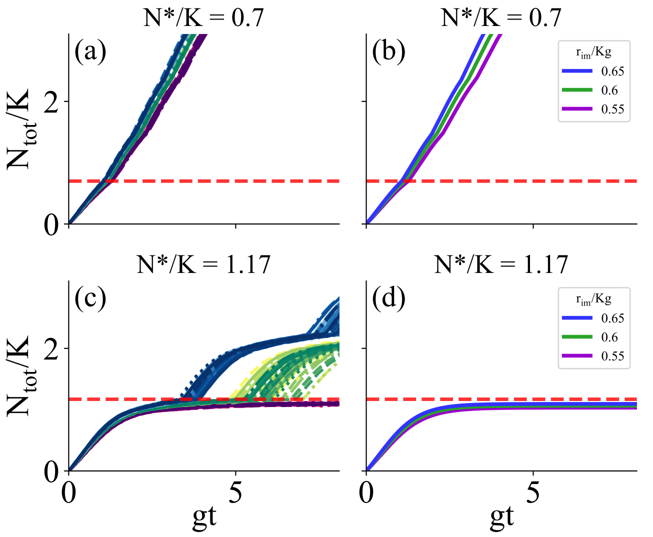


FIG. 4. Comparison between stochastic (left-hand panels) and deterministic (right-hand panels) biofilm growth trajectories. (a,b) Parameters in the deterministic growth regime; $N^*/K = 0.7$. (c,d) Parameters in the stochastic growth regime; $N^*/K = 1.17$. In all cases, $r_{\rm mig}/g = 0.8$, $r_{\rm det}/g = 0.5$, $K$=10,000 and the immigration rate is varied: $r_{\rm im}/gK$= 0.55, 0.6 and 0.65. The red dashed lines indicate the biofilm transition threshold $N^*/K$.
